# Origin of Protein Quake: Energy Waves Conducted by a Precise Mechanical Machine

**DOI:** 10.1101/2022.04.12.488080

**Authors:** Huiyu Li, Shanshan Wu, Ao Ma

**Affiliations:** Richard Loan and Hill Department of Biomedical Engineering, The University of Illinois at Chicago, 851 South Morgan Street, Chicago, IL 60607

## Abstract

A long-standing challenge in protein biophysics is to understand protein quake in myoglobin—the structural dynamics responsible for redistributing the excess heme energy after photolysis. Despite extensive efforts, the molecular mechanism of this process remains elusive. Using the energy flow theory, we uncovered a fundamental new phenomenon: the heme energy is redistributed by energy waves with a ubiquitous fundamental frequency and two overtones. Energy waves emanate from the heme into the myoglobin backbone via a conduit of five consecutive dihedrals of the proximal histidine, then travel quickly along the backbone to reach sidechains across the protein. This mechanism is far more effective than the diffusion-based mechanism from previous studies because waves are systematic while diffusion is random. To propagate energy waves, coordinates must cooperate, resulting in collective modes that are singular vectors of the generalized work functional. These modes show task partitioning: a handful of high-energy modes generate large-scale breathing motion, which loosens up the protein matrix to enable hundreds of low-energy vibrational modes for energy transduction.

---

Proteins are the building blocks of biological systems responsible for most biological functions. Understanding the mechanism of protein function is of paramount importance. The central dogma of protein science is that structure determines function. The critical link between structure and function is the functional dynamics (*1, 2*) that carries out protein function: structure determines function because the specific structure enables the desired functional dynamics. Understanding protein functions requires understanding the detailed molecular mechanism of functional dynamics.

The physical picture behind functional dynamics is that a protein has a multitude of functional structures, corresponding to basins in the underlying energy landscape that control the behavior of the protein (*3*). Transitions between these functional structures, which are carried out by functional dynamics, are often required for protein function. Following the pioneering work by Frauenfelder and co-workers (*2*), this idea has played a central role in experimental and computational studies of protein biophysics for the past few decades (*4-8*).

One of the first and most studied functional dynamics is the protein quake, a concept derived from pioneering studies of the structural dynamics of myoglobin (**Mb**) after photolysis by Ansari et al (*1*). The physical picture of protein quake is that the deposition of excess energy at a local site causes a strain in a protein, which then propagates through the protein matrix like an earthquake propagating through the mantle of earth. There are two essential hypotheses in this immensely appealing picture: 1) wave-like energy transduction through the protein matrix, 2) quake-like structural dynamics. Both ideas have attracted long-lasting and recurring interests and efforts from experimentalists and theorists alike.

The energy transduction during protein quake has been the subject of many spectroscopic studies since the pioneering work by Hochstrasser and co-workers in 1978 (*9-16*). These experiments were complemented by a series of computational studies dating back to the work by Henry et al in 1986 (*17-19*). The general conclusion is that the excess heme energy due to the absorption of a photon will be dissipated in ∼5 ps through anisotropic diffusion, suggested by computational studies by Straub and co-workers (*19, 20*). However, as originally pointed out by Lian et al (*16*), the slow time scale of diffusion contradicts the observed fast energy transfer, leading to the hypothesis of ballistic energy transfer by collective modes. The open question is: What are these modes?

On the structural dynamics side, recent advancements in X-ray free electron lasers (**X-FEL**) made it possible to “film” molecular movies of ultrafast protein dynamics (*21*), kindling the efforts to capture the protein quake in action (*22*). Time-resolved femtosecond crystallography, complemented by QM/MM simulations, showed collective response of the protein within 500 fs of photolysis of Mb that originates from the coupling of heme vibrations to global protein modes (*23*), but the specific nature of the protein modes is unclear. Similarly, time-resolved solution phase small- (**SAXS**) and wide-angle X-ray scattering (**WAXS**) showed an increase of the radius of gyration (*R*_*g*_) of Mb within 1 ps, followed by damped oscillations with a period of ∼3.6 ps (*24*). Another study using time-resolved SAXS and WAXS detected a single pressure peak propagating through the protein within 500 fs (*25*).

Despite many years of efforts and the abundance of information available on both aspects of protein quake, some critical links are still missing: 1) Is energy transduction wave-like as originally conjectured or diffusion-like as suggested by computational studies? 2) What is the molecular pathway for the energy transduction and what is the role of each residue or coordinate on this pathway? 3) What are the collective modes responsible for the energy transduction during a protein quake? These are the questions we strive to answer in the current paper.

We focused on the prototype of protein quake: the energy transduction and structural dynamics of Mb after photolysis. Previous computational studies were built on the assumption that this process is dominated by diffusion and focused on delineating the microscopic models for the friction kernel, which is the determining factor for diffusion (*19, 26-28*). In contrast, we did not make any model assumption, but focused on exactly accounting for every change in the system’s energy and ensuring there is no miscount of any factor. Our analyses are based on the recently developed energy flow theory, which allows us to rigorously map out the energy flow of each coordinate, and generalized work functional, which allows us to determine the cooperativity between coordinates (*29-31*).

Contrary to the diffusion-centric mechanism from previous studies (*17, 19*), we found that the excess heme energy is redistributed by energy waves with a ubiquitous fundamental frequency and two overtones. The excess heme energy is channeled into the Mb backbone by the proximal histidine (**His93**) that directly bonds with heme iron and then dispatched across the protein matrix. To propagate the energy waves, different coordinates must cooperate closely and Mb acts like a precise mechanical machine. The mechanism of this machine is best described by the singular vectors of the generalized work functional. Contrary to diffusion, waves are directional and systematic, thus they are more effective in distributing the heme energy. While diffusion-centered mechanism suggests that redistribution of heme energy resembles ink spreading in still water, the mechanism we found alludes to tiny objects ridding on ripples. The latter travels much faster than the former.

## Results

To simulate the protein quake of Mb after photolysis, extra kinetic energy is deposited to heme atoms *t* = 0 to mimic the excess energy from absorbing a photon. Afterwards, the system is simulated using constant energy molecular dynamics simulation (details in Methods).

### Energy flow theory

Our analysis is based on the potential energy flow (**PEF**) of individual coordinates (details in Supplementary Information (**SI**) and refs. (*29, 30*)). The PEF of a coordinate *q*_*i*_ along a dynamic trajectory *α* is the mechanical work done on *q*_*i*_ (*29*):

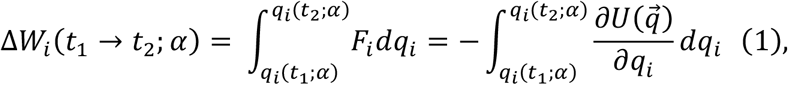

Where 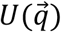 is the potential energy of the system, 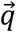 is the position vector of the system in the configuration space, and *F*_*i*_ is the total force, including forces from both solvents and the protein, exerted on *q*_*i*_. Because Δ*W*_*i*_(*t*_1_ → *t*_2;_ *α*) is the change in 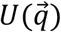 caused by the motion of *q*_*i*_ alone, it is the energy cost of the motion of *q*_*i*_. A change in 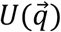 can be exactly decomposed as: Δ*U*(*t*_1_ → *t*_2_; *α*) = − ∑_*i*_Δ*W*_*i*_(*t*_1_ → *t*_2_; *α*), where the summation is over all the coordinates. This ensures that there is no missed count or overcount of any factor in the PEF of each coordinate.

To understand the mechanism of a protein quake, we compute PEFs averaged over an ensemble of trajectories, each trajectory simulates an instance of protein quake:

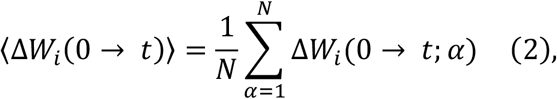

where *t* = 0 is set to be the time of photolysis, when the excess kinetic energy is deposited to the heme, *N* is the number of trajectories in the ensemble.

### Energy transduction by waves

The PEFs of an internal coordinate along different trajectories show distinct patterns (Fig. S1). Consequently, for each coordinate *q*_*i*_, we used k-means clustering (details in SI) to divide 8,000 trajectories into six ensembles *E*_*s*_(*q*_*i*_); *s* ∈ {0, …, 5}. The PEFs along trajectories in each ensemble resemble one pattern. For each coordinate *q*_*i*_, *E*_0._(*q*_*i*_) ≃ *E*_1_(*q*_*i*_) ≃ 3,300, *E*_2_ (*q*_*i*_) ≃ *E*_3_(*q*_*i*_) ≃ 540, *E*_4_(*q*_*i*_) ≃ *E*_5_(*q*_*i*_) ≃ 160. For two different coordinates *q*_*i*_and *q*_*k*_, *E*_*s*_ (*q*_*i*_) ≠ *E*_*s*_ (*q*_*k*_). Moreover, *E*_*k*_ (*q*_*i*_) ∩ *E*_*r*_(*q*_*k*_) ≠ 0; *r, s* ∈ {0, …, 5}; *r* ≠ *s*; its value indicates how strongly the motions of *q*_*i*_ and *q*_*k*_ are correlated.

Figure 1a shows the average PEFs of a backbone dihedral for all its 6 ensembles. Strikingly, the PEFs are wave-like. The dashed lines are the best fit of each PEF to the functional form *A* sin(*ωt* + *nπ*) ; *n* = 0, 1. The fits are nearly perfect, confirming that the PEFs are sine waves, with the 6 ensembles corresponding to 0.25, 0.75 and 1.25 period and initial phase of 0 and *π*, respectively. The fitted parameters (Table S1) correspond to a fundamental frequency of 0.32 ± 0.01*ps*^−1^ and its third and fifth overtones.

**Figure 1:**
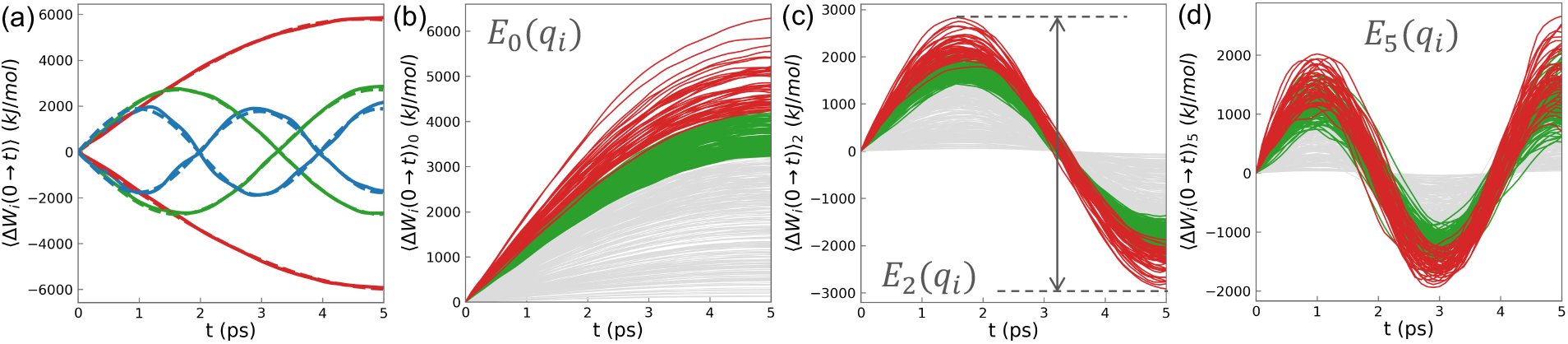
PEFs of backbone dihedrals are sine waves. (**a**) PEF of a backbone dihedral averaged over all the six ensembles. Dashed lines are the best fits to *A*_*t*_ *si*n (*ω*_*i*_*t* + *ϕ*_*i*_). (**b** to **d**) PEFs of all the backbone dihedrals averaged over *E*_0_(*q*_*i*_),*E*_2_(*q*_*i*_) and *E*_5_(*q*_*i*_), respectively.

Figure 1 shows the average PEFs for all the backbone dihedrals for *E*_0._(*q*_*i*_), *E*_2_(*q*_*i*_), *E*_4_(*q*_*i*_). All the PEFs are sine waves with essentially the same fundamental frequency. Each internal coordinate has 6 different excitation modes, corresponding to different combinations of overtones (1, 3 or 5) and phases (0 or *π*). In a single trajectory, different coordinates adopt different excitation modes; the initial configuration and momenta of Mb determines which excitation mode each coordinate adopts. These results support the idea that a protein quake uses waves for energy transduction, as Ansari et al originally conjectured in 1985 (*1*)!

### Energy level of individual coordinates

Each coordinate appears to have an intrinsic capacity for PEF that is independent of its mode of excitation. For a coordinate *q*_*i*_, the maximum of ⟭Δ*W*_*i*_(0 → *t*)⟩_0_ is much higher than the maximum of ⟨Δ*W*_*i*_(0 → *t*)⟩_2_, but ⟨Δ*W*_*i*_(0 → 5 *ps*)⟩_0_ = ⟨Δ*W*_*i*_(1.5 *ps* → 5 *ps*)⟩_2_ = −⟨Δ*W*_*i*_(0 → 5 *ps*)⟩_1_ = −⟨Δ*W*_*i*_(1.5 *ps* → 5 *ps*)⟩_3_ (gray arrow in Fig. 1c). This is true for all the coordinates. This intrinsic PEF capacity of a coordinate can be considered its energy level.

### Molecular pathway and mechanical mechanism of energy transduction in Mb

Figure 2 shows the PEFs of all the other types of internal coordinates averaged over their own *E*_0_ (*q*_*i*_). There are two essential features. First, PEFs of different types of coordinates differ significantly and there is a clear hierarchy. The first tier consists of dihedrals and bond angles of the backbone; their maximum PEFs can reach several thousand kJ/mol. The second tier consists of dihedrals, bond angles and improper dihedrals of the heme and His93. Their maximum PEFs can reach 1400 kJ/mol. The third tier consists of the backbone improper dihedrals, and dihedrals, bond angles, improper dihedrals and bonds of sidechains. Their maximum PEFs are well below 300 kJ/mol. In particular, the PEFs of backbone and sidechain bonds are extremely low—below 80 kJ/mol. Clearly, different types of internal coordinates serve distinct functional roles.

**Figure 2:**
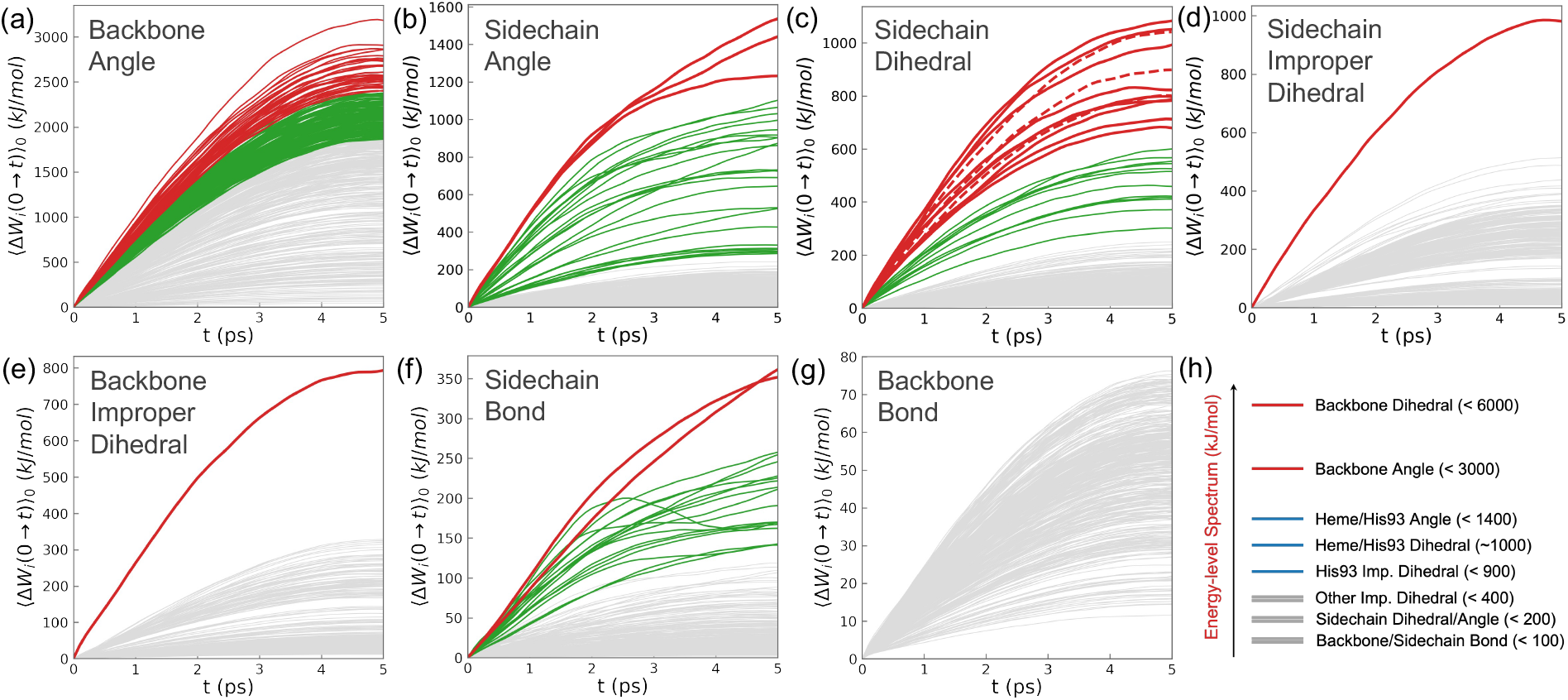
PEFs of different types of internal coordinates averaged over *E*_0_(*q*_*i*_). (**a** to **g**) PEFs of different types of internal coordinates, colored based on their magnitudes. In (**b, c, f**), all the red and green curves are PEFs of bond angles, dihedrals and bonds of heme or His93.. In (**d, e**), the two red curves are PEFs of improper dihedrals *δ*_1_, *δ*_2_ of His93 shown in Fig. 3d. (**h**) A schematic showing the hierarchy of PEFs of different types of internal coordinates.

Second, the main channel for redistributing the excess heme energy is through His93 into the protein backbone. Here, heme is the energy source, protein backbone is the repository, and His93 is the connector in between. This connector consists of two junctions, one with the heme and the other with the protein backbone, and a duct in between. The duct consists of three dihedrals *𝒳*_1_, *𝒳*_2_, *𝒳*_3_, and the two junctions are improper dihedrals *δ*_1_ and *δ*_2_ (Fig. 3). Both a *δ*_1_ and *δ*_2_ have PEF >800 kJ/mol (Fig. 2d,e), far above the maximum PEFs of all the other backbone and sidechain improper dihedrals, which are below 300 kJ/mol. Basically, the excess heme energy is conducted into His93 via the vibrations of *δ*_2_, then propagates through His93 sidechain via the puckering of the imidazole ring, controlled by *𝒳*_1_, *𝒳*_2_ and *𝒳*_3_, before passing into the backbone via the vibrations of *δ*_1_. The two junctions utilize improper dihedrals instead of other types of internal coordinates because their vibrations require little overall motion of the heme and His93 sidechain, which are difficult in the crowded environment of the heme pocket. The duct utilizes dihedrals because large-amplitude motion of the imidazole ring is required to convey high energy.

**Figure 3:**
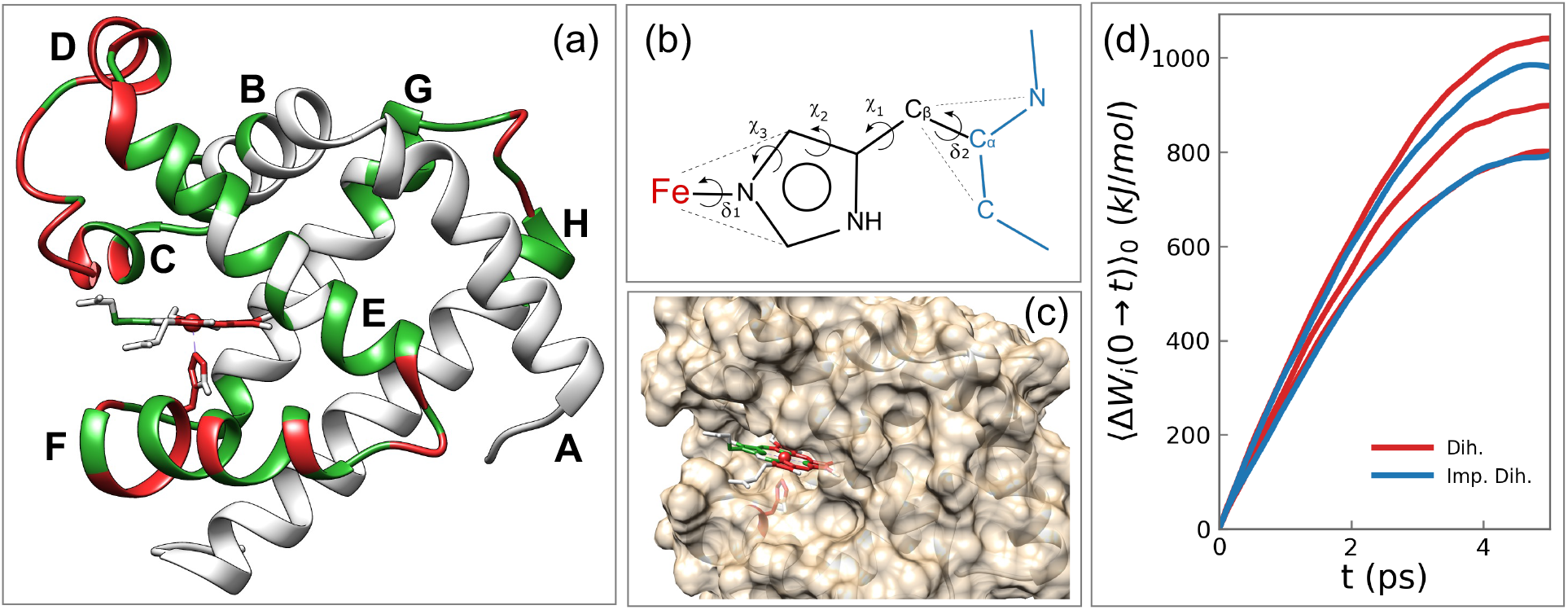
Molecular pathway for energy transduction in Mb. **(a)** Structure of Mb with different coordinates colored based on the magnitudes of their PEFs. **(b)** His93 as the connector between the heme and the protein backbone (atoms in **blue** color). The five internal coordinates that are critical for channeling the heme energy are labeled. **(c)** A close look at the difference in PEFs through the parts of the heme facing the interior and exterior of Mb. **(d)** PEFs through *δ*_1_, *δ*_2_ (**blue**) and *𝒳*_1_, *𝒳*_2_, *𝒳*_3_ (**red**).

To visualize the molecular pathway for energy transduction, we divide the internal coordinates into groups based on the magnitudes of their PEFs and colored them accordingly (Fig. 2). The groupings are based on comparing PEFs of coordinates of the same type (e.g. we compare PEFs of backbone dihedrals with each other, but do not compare PEFs of dihedrals to PEFs of bond angles). In the Mb structure in Fig. 3a, all the atoms of an internal coordinate are in the same color as its PEF. Therefore, the colors in Fig. 3a show the magnitudes of PEFs through different parts of Mb. The part of heme facing the interior of the heme pocket is “red hot” while the part facing the solvents has much lower PEFs (Fig. 3c). This suggests that heme-protein interactions are much more important for dispensing the heme energy than heme-water interactions, a conclusion different from previous computational studies (*19*). The entire His93 side chain is red because it channels the heme energy into helix F, which then propagates along two pathways. The major pathway heads towards the turn between helices F and E and then moves up helix E towards the CD-turn, the part of Mb that experiences the highest PEFs. The minor pathway is towards the turn between helices F and G. Overall, loops connecting helices experience the highest PEFs, while long helices have low PEFs, with helices E and F experiencing higher PEFs than the other helices.

Together, these results suggest that the redistribution of the excess heme energy is carried out by a mechanical machine that employs a hierarchy of energy flow channels. This machine operates similarly to a railroad system: the heme is the central terminal, protein backbone is the railroad, sidechains are the train stations, energy waves are the trains and bits of the excess heme energy are the passengers. After the heme energy is conducted into the backbone through His93, it rides on the energy waves to travel quickly along the backbone. When it reaches a sidechain on the way, a fraction of it is dispatched into the sidechain, with the size of the fraction determined by the size of the sidechain. In this way, heme energy is quickly redistributed over the entire protein matrix. This mechanism is much more effective than diffusion because wave is a directional motion while diffusion is random.

### Energy waves require a structural dynamics with global cooperativity

Figure 4a shows the PEFs of all internal coordinates averaged over *E*_0_(*ϕ* − 124), where *ϕ* − 124 is the *ϕ* dihedral of Gly124. Aside from the PEF of *ϕ* − 124 itself, PEFs of all but four dihedrals are vanishingly small. This is because *E*_0_(*ϕ* − 124) ∩ *E*_*s*_(*q*_*i*_) ≃ *E*_0_(*ϕ* − 124) ∩ *E*_*s*+1_(*q*_*i*_); *s* = 0,2,4, leading to strong cancelations and vanishing PEFs. In contrast, for the four dihedrals *ψ, ω* - 124, *ψ* of Phe123, and *ω* of Ala125 (blue lines), *E*_0_(*ϕ* − 124) ∩ *E*_1_(*q*_*i*_) ≫ *E*_0_(*ϕ* − 124) ∩ *E*_*s*_ (*q*_*i*_) (*s* ∈ {0, 2, 3, 4, 5}). Therefore, their PEFs averaged over *E*_0_(*ϕ* − 124) are dominated by their corresponding ⟨Δ*W*_*i*_(0 → *t*)⟩_1_, leading to the high PEFs in Fig. 4a. This situation indicates strong correlations between the modes of excitation of *ϕ* − 124 and its four neighboring dihedrals. This type of correlation exists extensively between dihedrals of a residue *i* and dihedrals of residues *i* ± 1. These local correlations concatenate into a chain of correlations that extends through the entire protein backbone, leading to global cooperativity between internal coordinates.

**Figure 4:**
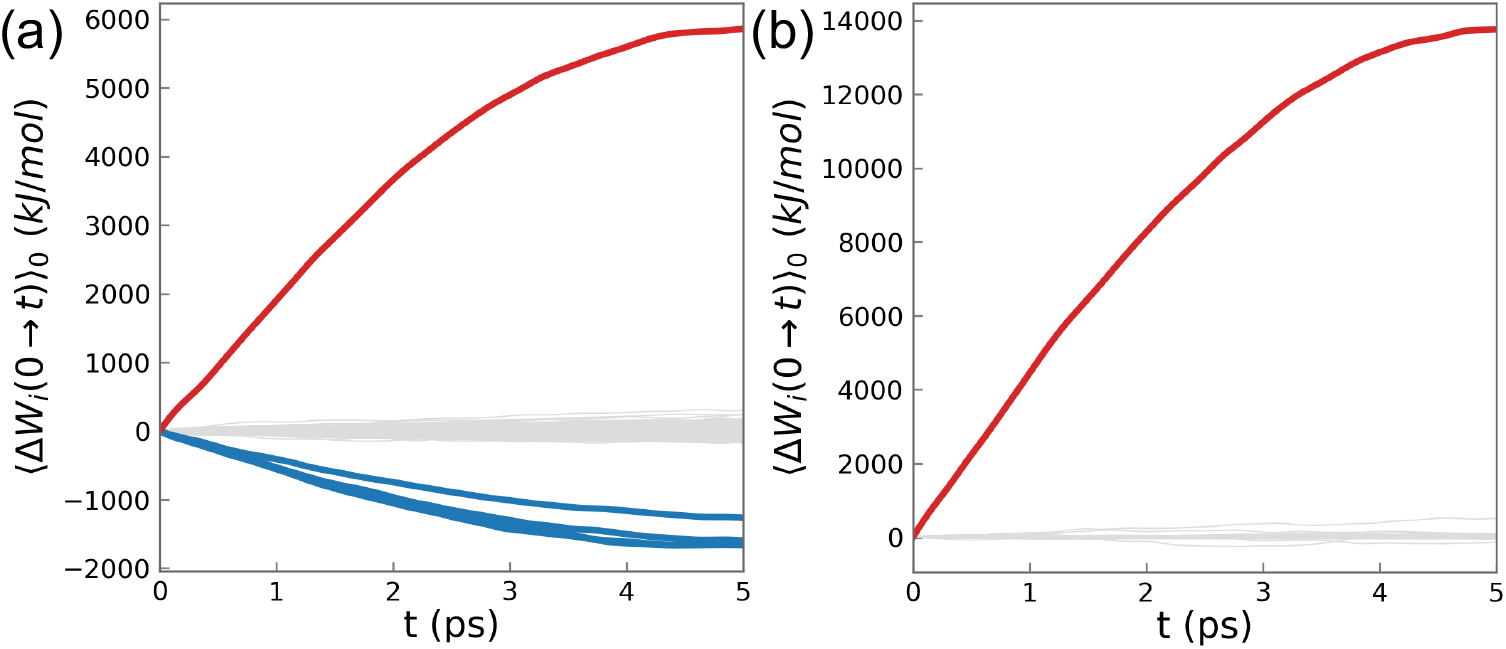
Correlations between different coordinates. **(a)** PEFs of all the internal coordinates averaged over *E*_0_ (*ϕ* − 124). **Red**: PEF of *ϕ* − 124; **Blue**: PEFs of *ω ψ* − 124, *ψ* − 123, *ω* − 125; **Gray**: PEFs of all the other dihedrals. **(b)** PEFs of all the singular coordinates averaged over *E*_0_(*u*_0_). **Red**: PEF of *u*_0;_ **Gray**: PEFs of all the other singular coordinates.

This global correlation is critical for function. While only 368 kJ/mol excess heme energy needs to be redistributed across Mb, PEFs of individual coordinates (Figs. 1, 2) far exceed this value because they are required for the systematic structural dynamics that is necessary for sustaining and propagating energy waves. The global correlation ensures that the high negative PEFs of one set of coordinates are balanced out by the high positive PEFs of another set of coordinates, keeping the overall energy cost of the systematic structural dynamics low. This enables redistributing the heme energy via energy waves.

### Generalized work functional

The existence of global correlation means the structural dynamics during a protein quake is best described by collective modes. To uncover these modes, we use the generalized work functional (*31*):

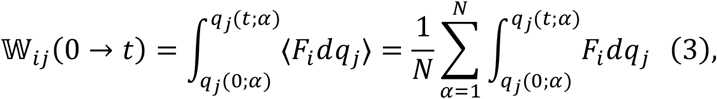

which summarizes the impact of *F*_*i*_ on the motion of *q*_*j*_. The collection of all the 𝕎_*ij*_(0 → *t*) is an asymmetric tensorial functional 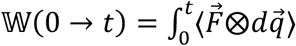 in the configuration space, as 𝕎_*ij*_ ≠ 𝕎_*ij*_. It summarizes the accumulated effects of the mechanical couplings between different coordinates as a function of the time since photolysis.

### Singular coordinates encapsulate the cooperativity

We use singular value decomposition to extract the left singular vectors of 𝕎(0 → *t*), which are the directions of forces that have the highest impacts on the motion of the system. Each singular vector *u*_*i*_ is a collective mode, which we refer to as a singular coordinate. Because 𝕎(0 → *t*) depends on time, the analytical expression of *u*_*i*_ is time dependent in principle. In practice, we found that expression of *u*_*i*_is time independent within numerical errors.

Figure 4b shows the PEFs of all the singular coordinates, constructed from the backbone dihedrals and all the coordinates of heme and His93, averaged over *E*_0_(*u*_0_). Unlike the case in Fig. 4a, only ⟨Δ*W*(*u*_0_)⟩ has high magnitude, while ⟨Δ*W*(*u*_*i*_)⟩ for all *i* ≠ 0. This means that the singular coordinates are uncorrelated, thus they serve functional purposes in independent manner. This is because the correlations between different internal coordinates are converted into the cooperativity between different components of *u*_*i*_, defined by the coefficients in the analytical expression of *u*_*i*_.

Figures 5 shows the PEFs of singular coordinates constructed from backbone dihedrals (Fig. 5a), bond angles (Fig. 5b), and the two together (Fig. 5c), aside from the coordinates of heme and His93. While the highest PEFs in the first two cases are far above, the highest PEFs of the third case are on par with the highest PEFs of backbone internal coordinates. The reason is that the PEFs of the bond angles compensate the PEFs of the dihedrals along every single trajectory, lowering the PEF of the singular coordinate. Figure 5d shows the dihedral and bond angle components of the velocity 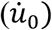 and force (*F*_0_) of 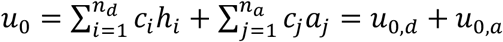_,:_ along a trajectory, where *n*_*d*_, *n*_*a*_ are the total number of backbone dihedrals and bond angles respectively, *h*_*i*_, *a*_*j*_ are individual dihedrals and bond angles, *c*_*i*_, *c*_*j*_ are the corresponding coefficients. Accordingly 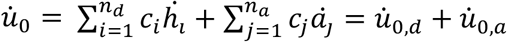, and 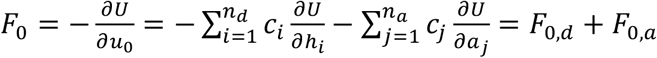 While *F*_0,*d*_ ≃ *F*_0,*a*_ along each trajectory (lower panel in Fig. 5d), 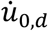 and 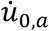 (upper panel) are always of opposite sign and similar magnitudes. Consequently, 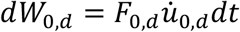 and 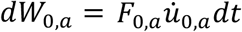 are of opposite sign and cancel with each other, significantly lowering 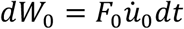. This precise energy compensation does not exist between individual dihedrals and bond angles; it only exists between the dihedral and bong angle components of the same singular coordinate (i.e. *u*_*i,d*_ and *u*_*i,a*_). This shows that singular coordinates precisely encapsulate the cooperativity between different coordinates, which is critical for the functional purpose of the protein quake. Therefore, singular coordinates can properly describe the structural dynamics during a protein quake.

**Figure 5:**
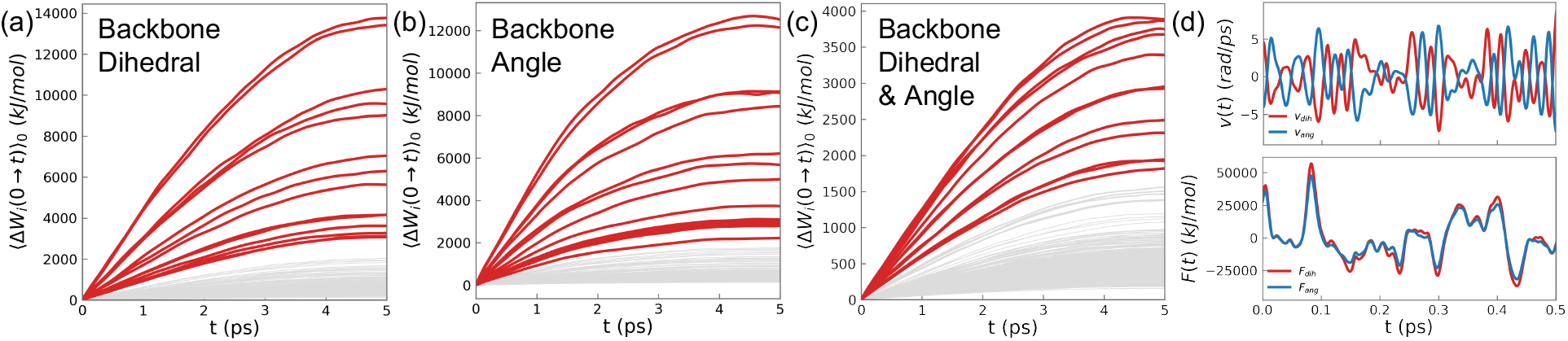
PEFs of singular coordinates. PEFs of all the singular coordinates constructed using all the internal coordinates of heme and His93 and backbone dihedrals **(a)**, backbone bond angles **(b)** and both **(c)**, averaged over *E*_0_(*u*_*i*_) **(d)** The dihedral (**red**) and bond angle (**blue**) components of the velocity (**upper**) and the force of *u*_0_ (**lower**) along a sample trajectory.

### Energy-level splitting

The PEFs of singular coordinates (Fig. 5) clearly separate into two groups. A small number of singular coordinates (13) have high PEFs and their “energy levels” are well separated. In contrast, all the other singular coordinates have low PEFs, which are densely packed into a narrow range. This clear difference in PEFs points to different functional roles. We refer to the first group (red lines in Fig. 5) as the functional modes and the second group (gray lines) as the dissipation modes. The clear separation in the “energy levels” upon transformation from internal to singular coordinates resembles the energy level splitting upon transformation from atomic to molecular orbitals.

### Task partitioning among singular coordinates

The clear separation between PEFs of the functional modes and the dissipation modes indicates that they serve distinct functional purposes. As shown in Fig. 6 and supplementary videos, a functional mode involves large-scale rigid body movements of the helices, with the amplitude proportional to its PEF. These movements are driven by small changes (up to 0.45°) in individual backbone dihedrals that have high PEFs (the red loops in Fig. 3a). They appear to be the breathing motion that was a central topic of Mb dynamics (*32*).

**Figure 6:**
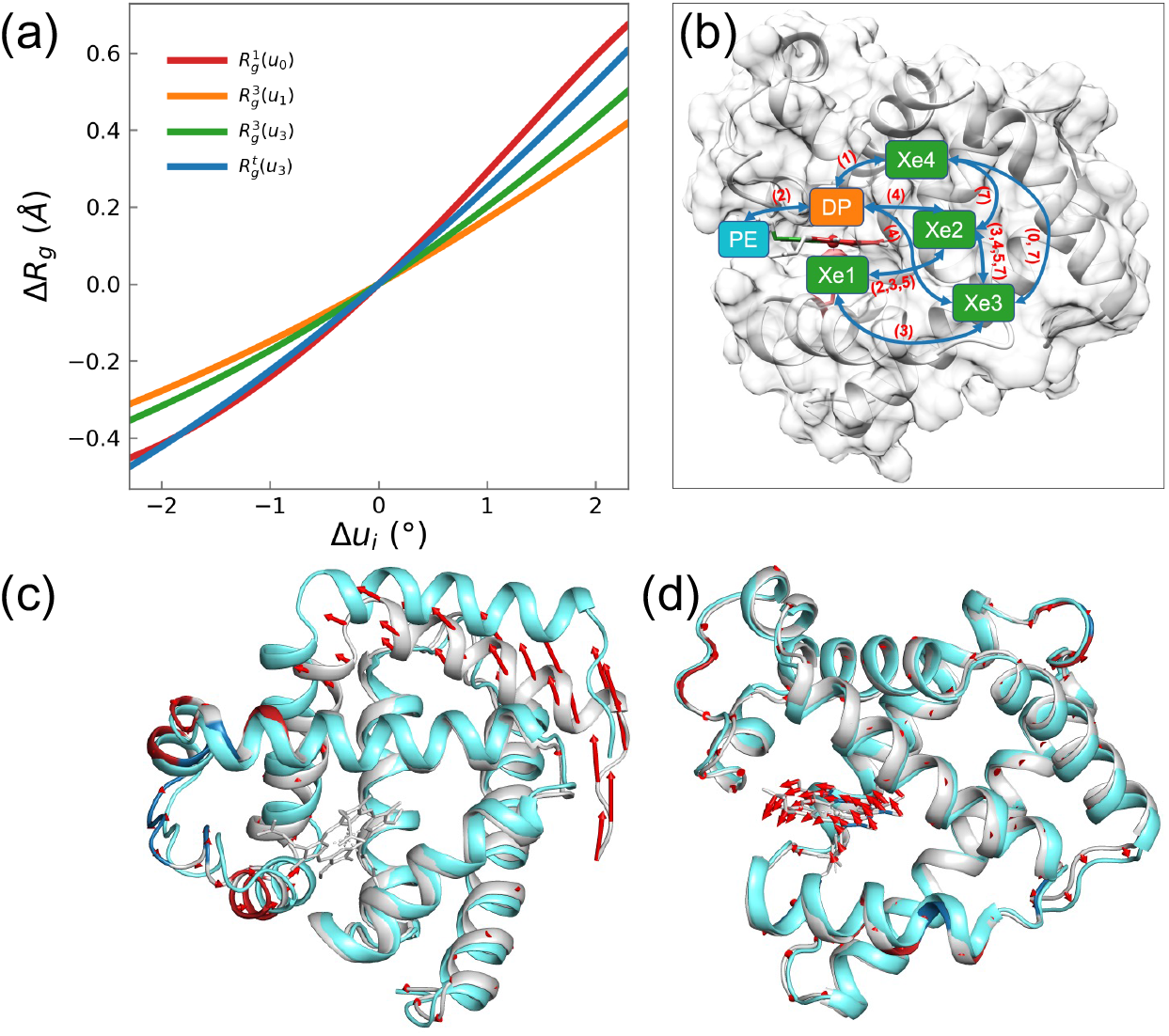
Effects of functional modes. (**a)** Change in *R*_*g*_ caused by displacements along a few functional modes 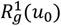 means change in the radius of gyration around the first principal axis of Mb caused by displacement along 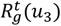 means change in the overall *R*_*g*_ **(b)** A schematic that summarizes the effects of displacements along different functional modes on the connectivity between internal cavities of Mb based on data in Table S2. A **blue** arrow connecting two cavities represents the channel between them. The numbers in parentheses are the indices of singular coordinates that control the open/close of the channel. **(c, d)** Direction of motion for singular coordinates *u*_0_ and *u*_41_. The crystal structure is colored **gray**; coordinates with coefficients > 0.1 or < − 0.1 are colored **red** (positive) and **blue** (negative). The structure after displacement along #_’_ is colored **cyan. Red** arrows indicate direction of displacement.

The breathing motion of Mb has two important effects: 1) change *R*_!_ of Mb, 2) control ligand migration through the internal cavities (distal pocket and xenon cavities) of Mb (*33*). Recent SAXS experiment by Levantino et al found an oscillation of *R*_*g*_ on a picosecond time scale after photolysis (*24*). We computed the change in *R*_*g*_ due to change in Mb structure induced by displacements along each functional mode. For each mode, we use the crystal structure *S*_*c*_ as the reference point and obtain new structures *S*_*i*_= *S*_*c*_ + Δ*u*_*i*_, with Δ*u*_*i*_= ±2.3°, and then calculate the *R*_*g*_ of *S*_*i*_. Figure 6a shows that functional modes *u*_0_ and *u*_3_ can change *R*_*g*_ on a similar scale as observed by Levantino et al.

Ligand migration has been the subject of many time-dependent Laue crystallography and computational studies (*32, 34-37*). However, it remains an open question what are the protein motions that control ligand migration between these cavities, which are disconnected from each other in the crystal structure. As shown in Fig. 6b and Table S2 (*38*), a functional mode can merge two or more cavities, enabling the ligand to migrate between these cavities. In this way, a functional mode acts as the gate to the channel connecting two or more cavities. Because motion along a functional mode can open or shut this gate, it regulates ligand migration process. In addition, functional modes can also modulate the volume of individual cavities, adding a further layer of control on ligand migration. Previous computational studies mapped the pathways of ligand migration, but they did not provide information on the specific motions that control these pathways. We showed how these pathways can be easily regulated by a small number of collective modes.

Another long-standing question on Mb dynamics is the coupling between heme vibrations and global motions of the protein matrix. Recent experiment on the ultrafast dynamics of Mb after photolysis using femtosecond crystallography strongly suggested existence of such coupling (*23*), but does not answer two critical questions: 1) What kind of global motions are coupled to heme vibrations? 2) How are they coupled together? To answer these questions, we identified singular coordinates in which heme dihedrals and bond angles with high PEFs (red curves in Fig. 2b, 2c) have the largest coefficients. These coordinates are dissipation modes with relative high PEFs (>1,000 kJ/mol). Among these four modes, *u*_18_ and *u*_19_ couples stretching motions of heme with small-amplitude rigid-body motion of all the long helices (i.e. helices A to H), driven by the loops connecting helices D and E, F and G, G and H. The coupling is mainly between the bond angles of heme and backbone dihedrals and bond angles of these loops, as shown in Fig. S2 and videos S3. On the other hand, *u*_39_ and *u*_41_ couple doming and twisting motions of the heme plane with rigid-body motion of helices A, B, E, F, driven by loops connecting C and D, E and F, G and H. The coupling is mainly between heme dihedrals and backbone dihedrals and bond angles of these loops and helix F (Fig. 6d). These modes are the representatives of the coupling between heme vibrations and global protein motions, as there are many other dissipation modes with smaller amplitudes that directly couple heme vibrations with backbone dihedrals and bond angles.

## Discussions

Compared to the large-scale conformational dynamics involved in ligand binding and allosteric transitions, the structural dynamics during a protein quake has small amplitude and is subtle. Even though it is fundamentally different from thermal fluctuations, it is a challenge to discern it from the latter. To uncover this intrinsic order buried under thermal noise, a precise method is critical. The energy flow theory allows us to map out the exact energy cost of the motion of every single coordinate during a protein quake, providing a comprehensive way to eliminate noise by proper ensemble average. This enabled us to uncover the detailed molecular mechanism of the energy transduction and structural dynamics of Mb induced by photolysis.

Contrary to the previous conclusion that this process is dominated by diffusion (*17, 19, 26*), we found that it is carried out by a well-organized hierarchy of energy flow channels synchronized by systematic structural dynamics that are encapsulated by the singular coordinates. The functional modes loosen up the protein to get the dissipation modes going so that energy waves can propagate. The excess energy of the heme is carried by this energy wave across the protein matrix and into the solvents. This mechanism is much more efficient than diffusion because waves are directional and systematic while diffusion is random.

Our results suggest that, in a functional process, the behavior of a protein is more like a fine-tuned mechanical machine designed for a specific purpose, rather than a stochastic system that relies on chance to achieve functional consequences. It is most tempting to speculate that functional processes utilize the same set of modes but excite them to different amplitudes. While energy or signal transduction utilizes synchronized energy waves through many modes, an activated process selectively excites one or a few functional modes to a very high level.

## Simulation Method

All simulations were performed using the molecular dynamics software suite GROMACS 2019.2 with CHARMM 36m force field (*39-41*). The sperm whale Mb molecule was solvated in a water box of 76 Å × 76 Å × 76Å. The simulation system consists of 41,267 atoms in total. The temperature was slowly increased to 300 K, followed by 40 ps of constant temperature molecular dynamics for equilibration (*19*). Afterwards, we run a long constant NVE MD simulation for 40 ns. From this trajectory, we select an initial condition every 5 ps as the starting condition for the subsequent photolysis simulations. To mimic the photolyzed state, an extra 368 kJ/mol of kinetic energy is deposited into the heme by rescaling the velocities of heme atoms, which amounts to increase the heme temperature by 400 K. More details on the energy flow calculations can be found in the SI.

## Acknowledgement

We thank the National Science Foundation for financial support (award: CHE-1665104).

## Notes

### Competing Interest Statement

The authors have declared no competing interest.

